# Structural and Biophysical Basis of the First DNA Aptamer Targeting the Dynamic p47 Adaptor of the p97 AAA+ Machine

**DOI:** 10.64898/2026.11.20.768169

**Authors:** Peter E. Kim, Yibo Liu, Matteo Pavlovic, Logan Fanning, Richard A. Manderville, Juewen Liu, Derek O’Flaherty, Rui Huang

## Abstract

Selective modulation of dynamic proteins within native multiprotein assemblies remains a major challenging for chemical biology. Here, we develop p47A1, the first DNA aptamer targeting the p97 adaptor p47 (NSFL1C) and uncover an unexpected allosteric mechanism of action. Target-immobilized SELEX identifies a highly convergent aptamer whose conformation is stabilized by Mg^2+^. Our comprehensive structural and biophysical characterization shows that p47A1 recognizes the SEP domain and adjacent SIM–SEP linker of p47 with low-micromolar affinity. Rather than simply occupying the p47 surface, aptamer binding remodels the intramolecular regulatory architecture of p47, disrupting intramolecular contacts and releasing the UBA domain into a highly dynamic state. Site-specific fluorescent labeling and mutational studies define the aptamer binding site. Importantly, reconstitution studies demonstrate that the aptamer engages p47 within the intact p47–p97 complex without disrupting the native complex or the basal p97 ATPase activity. Together, p47A1 provides a precise chemical biology probe for p47 and illustrates the potential of aptamers to target dynamic regulatory adaptors within native AAA+ protein assemblies.

## Introduction

Nucleic acid aptamers, single-stranded DNA or RNA oligonucleotides selected *in vitro* to recognize molecular targets with high affinities and specificities, represent a chemically versatile modality across chemical biology, molecular diagnostics, and targeted therapeutics.^1–4^ Compared to conventional antibodies, aptamers offer advantages such as modular chemical synthesis, minimal batch-to-batch variability, low immunogenicity, facile site-specific functionalization, and reversible thermal refolding.^1,5–7^ Their structural plasticity further enables recognition of shallow and extended protein interfaces that are challenging for conventional small molecules.^3,8^ Yet, despite more than three decades of SELEX (Systematic Evolution of Ligands by EXponential enrichment) discovery,^1,2^ the translational deployment of protein-targeting aptamers remains constrained by a fundamental bottleneck: the paucity of rigorous, residue-level structural, dynamic, and thermodynamic characterization and validation.^9–11^ Aptamers emerging from naïve selections frequently exhibit poorly defined binding parameters and conformational ensembles, non-specific electrostatic interactions, or diminished target engagement in the context of physiological multiprotein assemblies.^9,12–15^ Overcoming this barrier therefore demands an integrated strategy that combines aptamer selection with rigorous biophysical, structural and functional characterization and validation.

The AAA+ (ATPases Associated with diverse cellular Activities) molecular machine p97 (also known as valosin-containing protein, VCP) represents a compelling system in which such precision chemical probes are needed. This abundant homohexameric ATPase harnesses ATP hydrolysis to extract, unfold, and remodel polyubiquitinated proteins, thereby coordinating diverse proteostatic pathways including endoplasmic reticulum-associated degradation (ERAD), autophagy, endosomal sorting, chromatin regulation, and organelle membrane dynamics.^16–18^ Dysregulated p97 activity is implicated in human diseases. Pathogenic mutations are associated with multisystem proteinopathy type 1 (MSP1) and amyotrophic lateral sclerosis (ALS),^19,20^ whereas p97 overexpression is observed across multiple cancers and correlates with cancer aggressiveness and therapeutic resistance.^21–23^ Extensive medicinal chemistry efforts have focused on pharmacological inhibition of p97 using small molecules^21,24^ However, as an essential hub integrating multiple cellular pathways, global inhibition of p97 risks disrupting indispensable housekeeping functions and thereby narrowing the therapeutic window.

A conceptually attractive strategy for achieving greater pathway selectivity is to target the adaptor proteins that direct p97 toward defined cellular processes and substrate cohorts.^17,25^ Among the more than 30 known p97 adaptors, p47 (also known as NSFL1C) is particularly intriguing because it recruits p97 to mediate post-mitotic Golgi and endoplasmic reticulum membrane reassembly.^26–29^ This pathway is particularly active in proliferating and malignant cells, potentially creating a therapeutic window inaccessible to ATPase-directed inhibitors.^30–33^ Yet non-catalytic adaptors such as p47 pose substantial challenges for conventional small-molecule discovery. p47 comprises three folded domains connected by intrinsically disordered linkers, lacking deep hydrophobic pockets typically exploited by small molecules. To our knowledge, no synthetic small-molecule ligand or chemical probe has been reported for p47, and only one antibody fragment has been described to target its UBX domain.^25^

Here, we report the selection and comprehensive biophysical and structural characterization of p47A1, the first DNA aptamer developed to target p47. Target-immobilized SELEX yielded a highly converged aptamer that undergoes Mg^2+^-dependent conformational rearrangement and stabilization within its core. Integrated biophysical and NMR analyses reveal that p47A1 engages the SEP domain and adjacent SIM–SEP linker of p47 with low-micromolar affinity across various buffer conditions. Remarkably, p47A1 binding remodels the intramolecular regulatory architecture of p47, disrupting the intramolecular contacts and releasing the UBA domain into a highly dynamic state. Site-specific fluorescent labeling and mutational analyses further delineate the interacting region of the aptamer. Finally, we demonstrate that p47A1 engages p47 within the intact p47–p97 complex while preserving the native p47–p97 interface and the basal p97 ATPase activity. Together, this work establishes p47A1 as a precise chemical biology probe for allosteric modulation of p47 and demonstrates how aptamers can target dynamic regulatory interfaces within intact AAA+ protein assemblies.

## Results and Discussion

### Selection of p47-binding aptamers using SELEX

A target-immobilized selection strategy was employed to enrich DNA aptamers capable of binding p47. The p47 protein was covalently conjugated to NHS-activated agarose beads via primary amines on solvent exposed lysine residues, enabling stable presentation of the target during selection. A 73-nucleotide ssDNA library, consisting of a 30-nucleotide randomized region flanked by constant primer-binding sequences, was incubated with the immobilized protein. Following removal of unbound sequences through stringent washing, bound DNA was eluted and subsequently amplified by PCR to generate the input pool for subsequent rounds of selection (Figure 1A, Table S1). After ten rounds, the enriched pool was subjected to deep sequencing.

**Figure 1.**
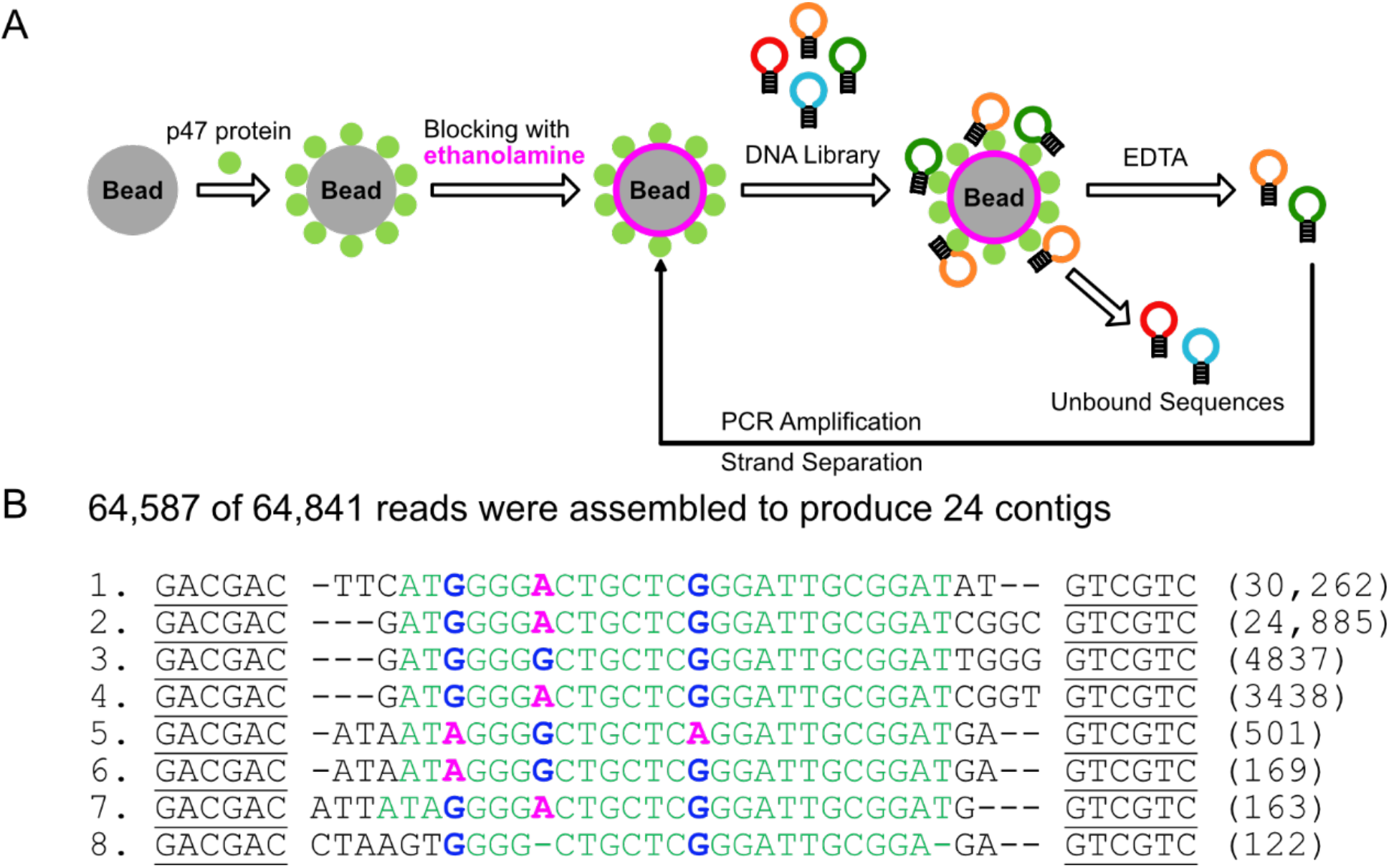
Selection of p47-binding aptamers via target immobilized-SELEX. (A) A schematic representation of the protein-immobilized aptamer selection procedure. (B) Alignment of the top 8 most abundant sequences out of a total of 64,841 sequences. The conserved sequences are shown in green. The underlined region represents the primer binding region that flanks the central randomized library region.

A total of 64,841 reads were obtained, of which 64,587 were classified into 24 sequence families using Geneious software, indicating a high degree of enrichment. The initial library was designed to adopt a hairpin-like architecture, which guided rational truncation of non-essential regions. Most of the primer-binding sequences were removed, while retaining six base pairs at each terminus to preserve structural stability, yielding minimized 42-nucleotide aptamers. Sequence analysis revealed strong convergence toward a dominant motif. The top eight most abundant sequences are shown in Figure 1B, with the two most enriched sequences accounting for approximately 85% of the total reads. Such pronounced enrichment suggests a highly favorable binding interaction between this motif and p47, likely reflecting both structural compatibility and favorable energetics. All enriched sequences contain a conserved core region (Figure 1B, green), with three positions exhibiting purine variability (G or A). The limited degeneracy suggests that these positions may tolerate substitutions without disrupting the overall fold, potentially maintaining key base-stacking interactions or preserving a binding-competent conformation rather than directly mediating specific contacts with the protein. Collectively, the high degree of sequence convergence, combined with the presence of a conserved structural motif and limited tolerated variability, supports the successful selection of aptamers that adopt a defined secondary structure optimized for p47 recognition. This convergence also suggests that the binding interface imposes strong structural constraints, favoring a narrow ensemble of functional sequences. We selected the most abundant sequence from the selection (Figure 1B, top) for further detailed characterization, which is hereafter referred to as p47A1 aptamer (Figure 2A).

**Figure 2.**
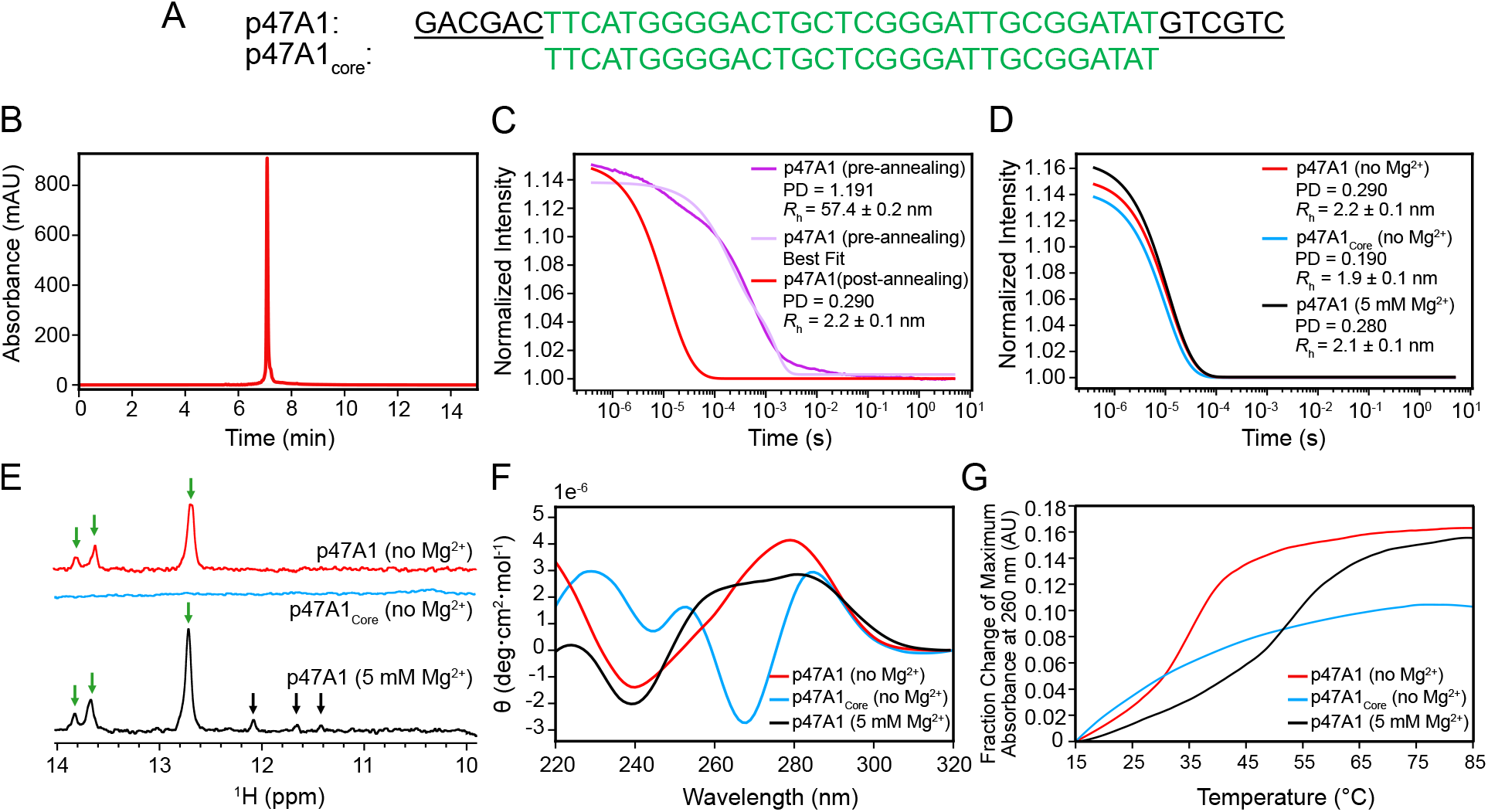
Biophysical characterization of the p47A1 aptamer. (A) Nucleotide sequences of the p47A1 aptamer (top) and the truncated variant, p47A1_core_, lacking the primer-flanking regions (bottom). (B) Anion exchange HPLC chromatogram of purified p47A1. (C) DLS autocorrelation functions of 120 μM pre-annealed (purple) and post-annealed p47A1 (red). (D) DLS autocorrelation functions of 120 μM p47A1 in the absence (red) and presence (black) of 5 mM MgCl₂, and p47A1_core_ in the absence of MgCl₂ (cyan). Fitted parameters for DLS are summarized in Table S3. Imino proton region of 1D ^1^H NMR spectra (E), CD spectra (F) and thermal denaturation profiles (G) of p47A1 in the absence and presence of 5 mM MgCl₂ were shown in red and black, respectively, while those of p47A1_core_ in the absence of MgCl₂ were shown in cyan. Measurements were performed in a buffer containing 10 mM Na_2_HPO_4_, 1.8 mM KH_2_PO_4_ (pH 6.6), 25 mM NaCl, and 5 mM MgCl₂ (wherever indicated).

### p47A1 core region is largely unstructured and undergoes Mg^2+^-induced conformational rearrangement and stabilization

We synthesized the p47A1 aptamer via solid-phase oligonucleotide synthesis and purified it using strong anion exchange chromatography for the following studies. The high purify (> 90%) of the aptamer is confirmed by the single, sharp symmetric peak in the HPLC analysis (Figure 2B). We then used dynamic light scattering (DLS) to characterize the hydrodynamic radius and structural homogeneity of the aptamer. Prior to thermal annealing, the sample exhibited high polydispersity (PD =1.191) and an average hydrodynamic radius (*R*_h_) of ∼ 57 nm (Figure 2C, purple and light purple). This hydrodynamic radius is significantly larger than that of the monomeric aptamer with a *R*_h_ of 2.1 nm as predicted by HYDROPRO^34^, implying the presence of higher-order oligomers or non-specific assembly post-synthesis. After thermal annealing, the aptamer transitioned to a monodispersed species with a substantially decreased *R*_h_ of 2.2 nm and a low polydispersity index (PD = 0.290) (Figure 2C, red), consistent with a structurally homogeneous and monomeric species.

To probe base-pairing and folding within the aptamer, we recorded 1D ^1^H NMR spectra focusing on the downfield imino proton region (10-14 ppm), which directly reports on hydrogen bonding. Prominent imino resonances, which correspond to stable base pairs, were observed at 12.7, 13.6, and 13.8 ppm (Figure 2E, red). These signals most likely originate from the complementary primer flanking regions (Figure 2A, underlined). This is confirmed by the absence of these resonances in the truncated variant, p47A1_core_, that lacks the primer-flanking region (Figure 2E, cyan; Figure 2A), suggesting no stable base-pairing in the core sequence of the aptamer (Figure 2A, green).

We then used circular dichroism (CD) spectroscopy to assess the secondary structure of the aptamer. The CD spectrum of the p47A1 aptamer exhibits characteristic features of a single-stranded DNA dominated by B-form base-stacking (Figure 2F, red), consistent with the presence of a hairpin closed by the flanking region.^35^ In contrast, p47A1_core_ is largely unstructured according to the CD spectra (Figure 2F, cyan). An estimated melting temperature of 36 °C for the p47A1 aptamer (Figure 2G, red), combined with the lack of characteristic thermal melting profile (Figure 2G, cyan) for p47A1_core_, further suggests a lack of stable base pairing in the core sequence of the aptamer beyond the terminal stem region.

Given the established role of Mg^2+^ in inducing structural, dynamic and oligomeric state changes in DNA,^36–38^ we next investigated the effect of Mg^2+^ on the oligomeric state and secondary structure of p47A1. Our results show that the hydrodynamic radius and homogeneity of the aptamer are not impacted by the addition of Mg^2+^ (Figure 2D, red *vs.* black). However, additional weak imino proton signals were observed at 12.1, 11.6, and 11.4 ppm in the 1D ^1^H NMR spectrum in the presence of Mg^2+^ (Figure 2E, black arrows), suggesting the formation of new base pairs. This finding is corroborated by a significant change in CD spectra upon the addition of Mg^2+^ (Figure 2F, red *vs.* black), which indicates conformational rearrangement and potential secondary structure formation within the core sequence of the aptamer (Figure S1). The stabilization of the secondary structure by Mg^2+^ is further evidenced by a substantial increase in the melting temperature (*T*_m_) of p47A1 from 36 °C to 52 °C in the presence of Mg^2+^ (Figure 2G, red *vs.* black).

### NMR characterization identifies the SEP domain and the SIM-SEP linker region of p47 as the primary binding sites for p47A1

To elucidate the structural determinants governing the interaction between the p47A1 aptamer and p47, we employed 2D NMR experiments to map the binding site of the aptamer on p47. p47 consists of three folded domains – the UBA, SEP and UBX domains, connected by flexible linkers where several functional linear motifs (*i.e.*, SIM, SHP_N_, and SHP_c_) were located (Figure 3A)^39,40^. Figure 3B shows the ^1^H-^15^N HSQC spectra of the full-length p47 in the absence and presence of one molar equivalent of the p47A1 aptamer. Binding of the aptamer induced significant chemical shift perturbations (CSPs) that localized primarily in the central region of p47, encompassing the SEP domain, the SIM motif, and the linker between SIM and SEP (Figure 3C). These regions also show severe intensity attenuation upon complex formation (Figure 3D), confirming direction interactions with the aptamer. Concurrently, a uniform decrease in peak intensity was observed for residues in the UBX domain with minimal chemical shift perturbations (Figure 3C&D), which we attribute to an increase in the rotational correlation time of the domain resulting from the increased size of the complex. Intriguingly, residues in the UBA domain and on the UBA-SIM linker displayed an increase in peak intensity upon aptamer binding (Figure 3D). Given our previous demonstration of transient intramolecular interactions between the UBA and SEP domains, this unexpected intensity increase in UBA suggests a relief of conformational exchange broadening^40^. Competitive binding of the p47A1 aptamer to the SEP domain might effectively disrupt the transient intramolecular interactions, liberating the UBA domain and its flanking linker and thereby increasing their local flexibility and backbone dynamics.

**Figure 3.**
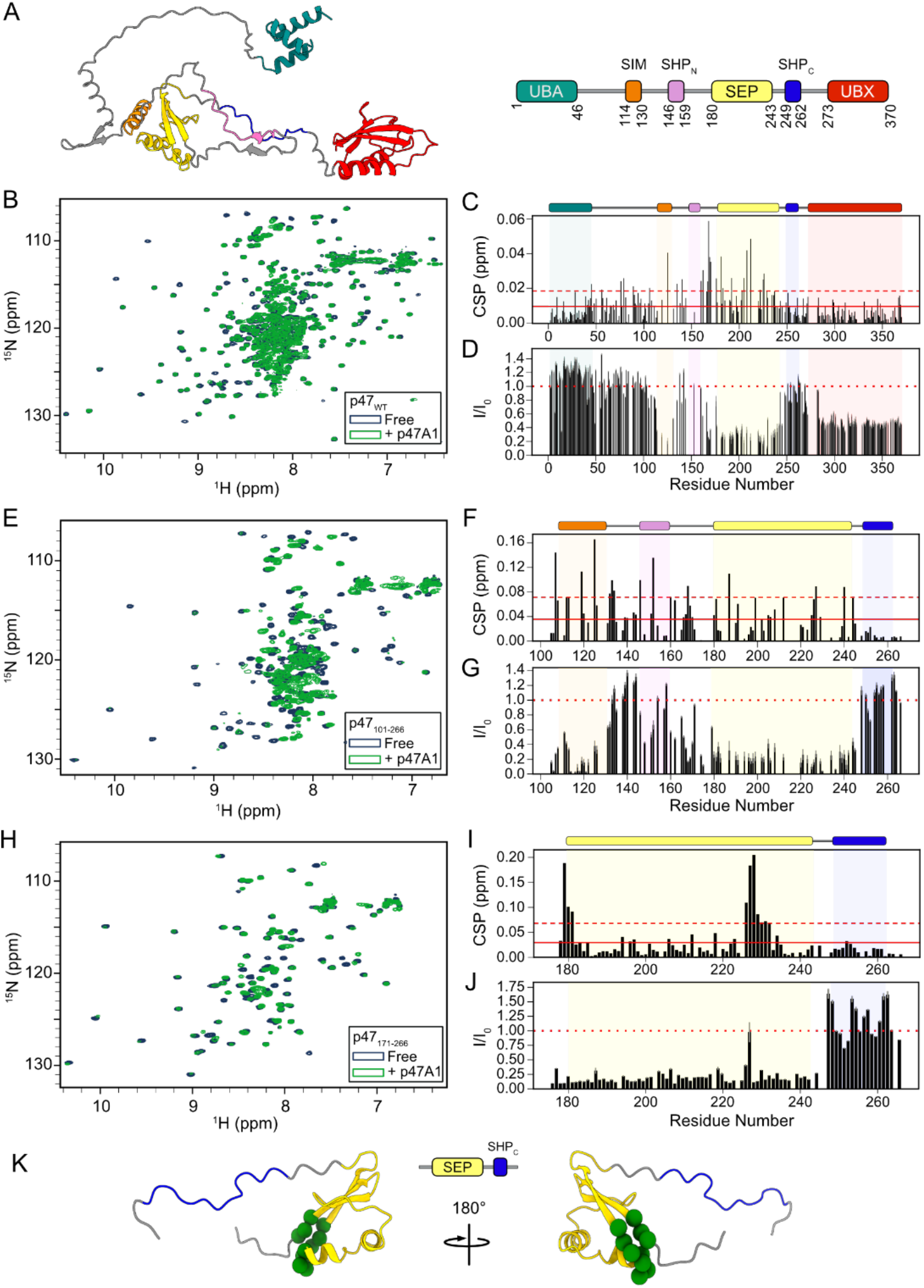
Identification of the p47A1 aptamer-binding sites on p47 by NMR. (A) Structural model of p47 predicated by AlphaFold 3 (left)^41^ and schematic of its domain organization (right). Superposition of 2D ^1^H–^15^N HSQC spectra of [U-^15^N] p47_WT_ (B), p47_101-266_ (E), and p47_171-266_ (H) in the absence and presence of the p47A1 aptamer at a 1:1 molar ratio. Chemical shift perturbations (CSPs) of p47_WT_ (C), p47_101-266_ (F) and p47_171-266_ (I) and peak intensity ratios between the aptamer-bound and free states (*I*/*I*_0_) of p47_WT_ (D), p47_101-266_ (G) and p47_171-266_ (J) are plotted by residue numbers. The solid and dashed lines in (C), (F), and (I) indicate the average CSP value and one standard deviation above the mean, respectively. The dotted line in (D), (G) and (J) indicates no change in intensity (*I*/*I*₀ = 1). (K) Structural mapping of the aptamer binding epitope onto p47_171-266_. Residues exhibiting significant CSPs are highlighted in green spheres.

Due to the large molecular size of the complex (∼ 50 kDa), a significant portion of resonances in the core SEP domain, particularly residues 200-250, suffered extensive line broadening and spectral crowding leading to missing information in this region (Figure 3B-D). To alleviate spectral crowding, reduce line broadening, and confirm essential domains for aptamer recognition, we recorded the ^1^H-^15^N HSQC spectra of two truncated variants of p47 in the presence and absence of the p47A1 aptamer (Figure 3E&H). The first construct, p47_101-266_, comprising the SIM motif, the SEP domain, the SHP_C_ motif, and the linkers in between, while the second minimal construct, p47_171-266_, encompassing solely the SEP domain and the SHP_c_ motif. Binding of the p47A1 aptamer to p47_101-266_ resulted in significant CSPs across the entire construct except for the SHP_c_ motif (Figure 3F), along with extensive line broadening in the SIM and SEP regions (Figure 3G). This profile is consistent with our observations in full-length p47 (Figure 3B-D), demonstrating that the binding interface is preserved in this truncated variant. Addition of the aptamer into p47_171-266_, a further truncated construct, allows for residue-specific tracking of both CSPs and intensity changes for nearly all residues in the SEP domain due to relieved spectral crowding (Figure 3I and J). The largest chemical shift changes were clustered at residues D179, V180, H181 and a short segment spanning A226–V231 (Figure 3I). These residues form a well-defined and localized epitope on the SEP domain (Figure 3K, green spheres), and the residues span diverse physicochemical properties (Figure S2), supporting a structure-driven recognition mechanism. It’s worth noting, however, that this minimal interface on p47_171-266_ is likely incomplete due to the absence of the SIM motif and the SIM-SEP linker, both of which likely contribute to aptamer recognition dynamically in the longer constructs (Figure 3C-D and F-G). Notably, a consistent feature across all the three constructs (p47_WT,_ p47_101-266_ and p47_171-266_) was an increase in peak intensities in the SHP_C_ motif upon aptamer binding (Figure 3D, G, and J). This phenomenon is likely due to the disruption of the transient intramolecular interactions between the SHP_C_ motif and the SEP domain by the aptamer competitively binding to an overlapping interface on the SEP domain.^40^ This displacement liberates the SHP_C_ motif and increases its local flexibility in a manner analogous to the release of the N-terminal UBA domain observed in p47_WT_ (Figure 3D).

### Thermodynamic characterization reveals low-micromolar affinity binding between the p47A1 aptamer and p47 under various buffer conditions

Having established that the p47A1 aptamer recognizes p47 through specific interactions with the SIM and SEP regions, we then employed isothermal titration calorimetry (ITC) to quantify the thermodynamic parameters governing these interactions. The binding energetics between the p47A1 aptamer and full-length p47 (p47_WT_), as well as two truncated variants (p47_101–266_ and p47_171–266_) that were previously used in the NMR experiments, were systematically evaluated under three buffer conditions varying in the presence or absence of Mg^2+^ and salt concentration. Buffer A contains 25 mM Bis-Tris (pH 6.6), 25 mM NaCl, 1 mM EDTA, matching the low-salt buffer condition used in the NMR investigations, whereas Buffer B contains 20 mM MES (pH 6.2), 150 mM NaCl, 10 mM KCl, and 5 mM MgCl_2_, replicating the high-salt condition wherein SELEX was performed, whereas Buffer C has the identical composition as Buffer A except for the presence of 5 mM MgCl_2_. Representative thermograms and the corresponding binding isotherms are shown in Figure 4 and S3, while the extracted thermodynamic parameters derived from a standard N-equivalent sites binding model are summarized in Table S3.

**Figure 4.**
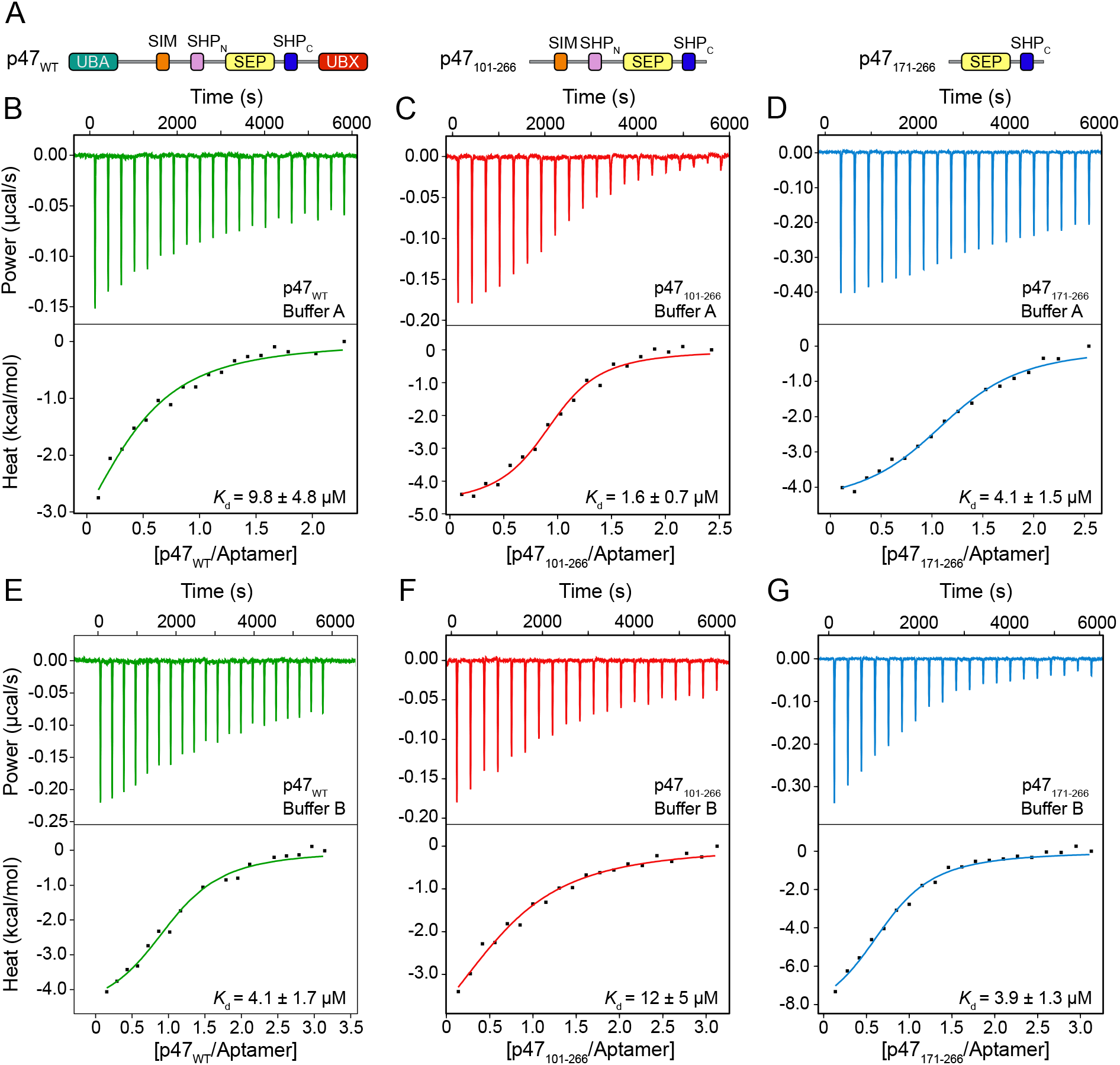
Binding energetics between the p47A1 aptamer and p47 variants by ITC. (A) Schematic representation of the full-length p47 (p47_WT_) (left) and the truncated variants, p47_101-266_ (middle), p47_171-266_ (right). (B–D) Representative raw ITC thermograms (top) and corresponding integrated binding isotherms (bottom) for titrations of p47_WT_ (B), p47_101–266_ (C), and p47_171-266_ (D) into the p47A1 aptamer in Buffer A (25 mM Tris, 25 mM NaCl, 1 mM EDTA, pH 6.6). (E–G) Representative raw ITC thermograms (top) and corresponding integrated binding isotherms (bottom) for titrations of p47_WT_ (E), p47_101–266_ (F), and p47_171-266_ (G) into the p47A1 aptamer in Buffer B (20 mM MES, 150 mM NaCl, 5 mM MgCl₂, and 10 mM KCl, pH 6.2). Solid lines in the binding isotherms represent the best fit to a two-state *N*-equivalent site binding model.

In Buffer A, the p47A1 aptamer binds p47_WT_, p47_101–266_, and p47_171–266_ with dissociation constants (*K*_d_) of 9.8 µM, 1.6 µM, and 4.1 µM, respectively (Figure 4B-D, Table S3). Notably, the aptamer exhibits the highest affinity to p47_101–266_ and the weakest affinity to full-length p47_WT_. This result strongly corroborates our NMR observations, which were performed under the same buffer condition (Figure 3). Our NMR results show that p47_101–266_ encompasses the complete binding interfaces to the aptamer, namely the SIM and SEP domains as well as the SIM-SEP linker (Figure 3C) but lacks the N-terminal UBA domain that engage in transient intramolecular interactions with the SEP domain and thus compete with aptamer binding (Figure 3D)^40^. Therefore, this variant is optimal for aptamer recognition, resulting in the highest binding affinity under this condition. Interestingly, unlike the cases for p47_101–266_ and p47_171–266_, the apparent binding stoichiometry between p47_WT_ and the aptamer deviates significantly from a 1:1 binding model (N = 0.50, Table S3), implying a more complex binding mode, likely originating from an intricate equilibrium between intermolecular aptamer binding and multiple transient intramolecular interactions within p47^40^, which often leads to a skewed stoichiometry value when fitted with a simplistic model^39^. Removal of the SIM and the adjacent SIM-SEP linker (p47_171–266_) results in a modest decrease in affinity from a *K*_d_ of 1.6 µM to 4.1 µM compared to p47_101–266_, confirming that besides the SEP domain, the SIM motif and SIM-SEP linker also contribute energetically to aptamer recognition and the stability of the complex, which is consistent with our NMR findings (Figure 3C&F).

Since Mg^2+^ coordination structurally stabilizes the p47A1 aptamer (Figure 2E-G), we then evaluated the effect of Mg^2+^ on the binding energetics between the aptamer and the p47 variants. Interestingly, the addition of Mg^2+^ (Buffer C) does not significantly alter the binding affinities of the aptamer to any of the p47 variants (Figure S3 and Table S3). We then characterized the interactions under a buffer condition with a higher ionic strength (Buffer B) (Figure 4E-G). Intriguingly, this condition causes a significant decrease in binding affinity for p47_101-266_ (1.6 µM *v.s.* 12 µM) while yielding no discernible change in *K*_d_ for p47_171-266._ This suggests that elevated ionic strength may selectively weaken key contacts involving the SIM motif or the SIM–SEP linker with the aptamer. This observation aligns well with the presence of positively charged stretch within the SIM–SEP linker adjacent to the SEP binding hotspots (Figure S4), which likely stabilizes aptamer-p47_101-266_ complex formation via favorable electrostatic interactions. Notably, unlike the apparent sub-stoichiometric binding observed under the low-salt condition (Buffer A, N = 0.50), the stoichiometry between the aptamer and p47_WT_ converges to unity with elevated ionic strength (Buffer B, N = 1.1). This shift suggests that higher salt concentrations may shield and destabilize the intramolecular contacts within p47_WT_, exposing the SIM-SEP core region for aptamer docking via a standard 1:1 binding mechanism.

Collectively, our NMR and ITC data demonstrate that the p47A1 aptamer recognizes an extended interface spanning the central region of p47, comprising the core SEP domain, the SIM motif, and the intervening SIM-SEP linker. Within this recognition epitope, the SEP domain serves as the primary anchoring site, whereas the SIM motif and the SIM-SEP linker provide auxiliary energetic contributions. Importantly, our findings reveal that the transient, multi-domain intramolecular network that defines the conformational ensemble of p47 can be strategically decoupled by modulating ionic strength or competitive aptamer binding.

### Identification of the core region in the p47A1 aptamer involved in protein recognition

Next, we sought to identify the core region in the p47A1 aptamer that is responsible for recognizing p47. Using the UNAfold Web Server^42^, the p47A1 aptamer was predicted to form two helical stems, one by base-pairing between the complementary 5′ and 3′ primer-binding regions (Figure 5A, Stem 1), and the other predicted to form between residues C17-G20 and G29-G32 (Figure 5A, Stem 2), partitioning the molecule into an internal loop and an apical loop (Figure 5A). Our previous 1D ^1^H NMR results confirmed that the terminal stem is stably folded (Figure 2E, green arrows), while the internal stem may be transiently formed and stabilized by Mg^2+^ (Figure 2E, black arrows). To determine whether the terminal stem (Stem 1) is directly involved in binding p47, we recorded 1D ^1^H NMR spectra of the aptamer variants in the absence and presence of p47. The p47_171-266_ construct was utilized in these experiments to optimize spectral quality and reduce relaxation-induced line broadening. Addition of p47_171-266_ to the p47A1 aptamer led to a notable reduction in the intensities of the imino proton signals corresponding to the terminal stem without inducing significant chemical shift perturbations of these signals (Figure 5B, green arrows, blue *v.s.* cyan). This is consistent with an overall increase in the global rotational correlation time upon complex formation. At the same time, new imino proton resonances emerge at 11.1, 11.6 and 12.1 ppm upon interaction (Figure 5B, black arrows, blue *v.s.* cyan), which directly indicates either binding-induced adaptive folding of the flexible regions of the aptamer or physical shielding of some imino protons from solvent exchange upon protein engagement. To confirm that these binding-induced structural changes originate from core region of the aptamer rather than the terminal stem, we investigated the truncated variant of the aptamer lacking the terminal stem, p47A1_core_ (Figure 5A). Addition of p47_171-266_ into the unstructured p47A1_core_ resulted in the emergence of the identical set of new imino proton peaks (Figure 5B, black arrows, red *v.s.* pink), supporting that the protein-induced folding or shielding event is localized within the core region of the aptamer. Furthermore, 2D ^1^H-^15^N HSQC spectra of ^15^N-labeled p47_101-266_ in the presence of either the full-length p47A1 aptamer or the truncated variant p47A1_core_ are superimposable (Figure 5C), indicating that the terminal primer binding stem is dispensable for protein recognition. This structural observation is further supported by the ITC results showing that deletion of the terminal stem resulted in no significant change in binding affinity towards p47_101-266_ (Figure S5 and Table S3).^38^

**Figure 5.**
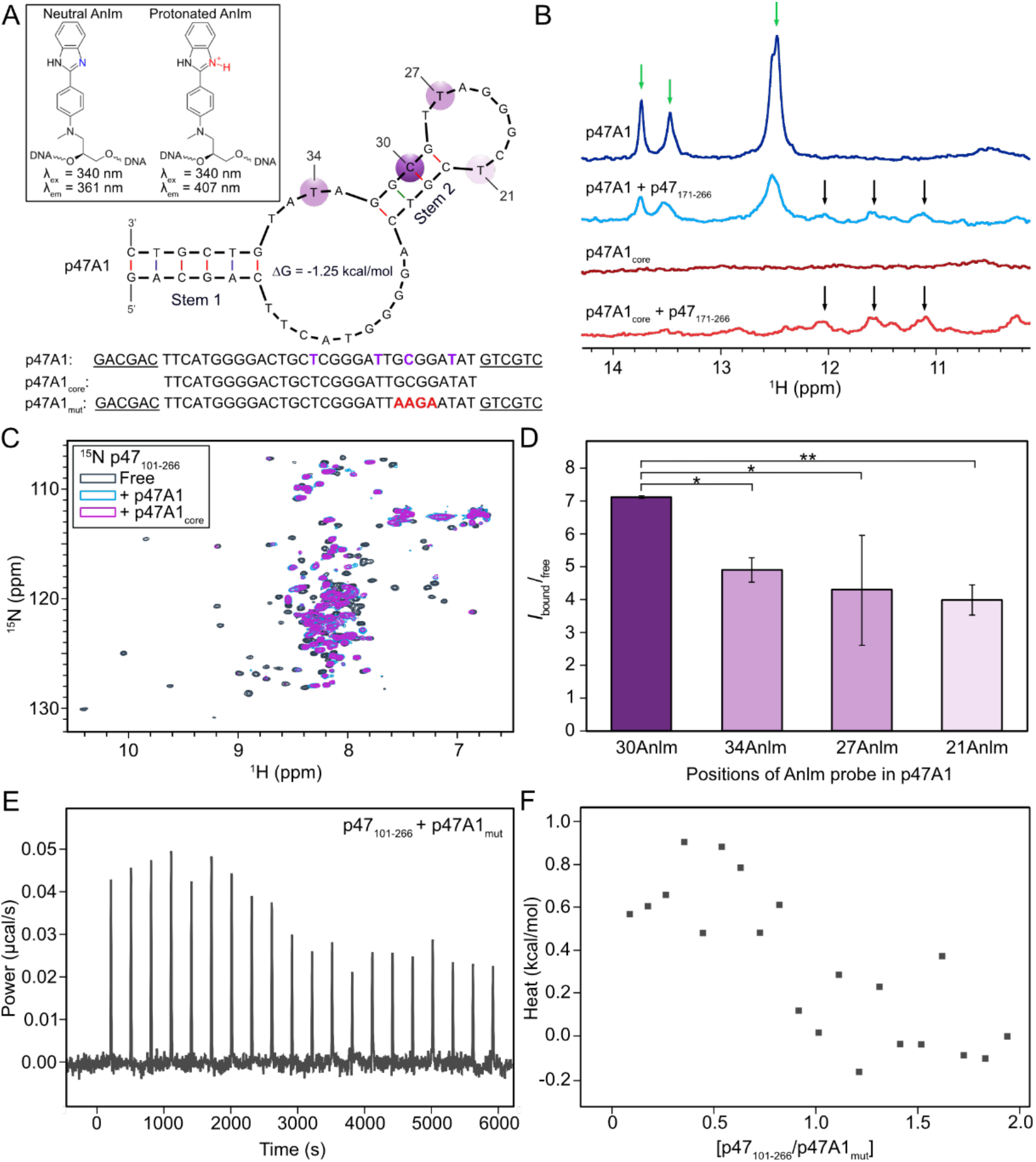
Identification of structural determinants in the p97A1 aptamer governing p47 recognition. (A) UNAfold-predicted secondary structure of p47A1.^42^ The Inset shows the chemical structures and photophysical properties of the AnIm photobase probe in neutral and protonated states. Sites of Anlm probe incorporation are highlighted with purple. Sequences of the aptamer constructs are shown below with the primer flanking regions underlined, Anlm labeling sites colored purple, and the 4-nt substitution highlighted in red. (B) The imino proton region of 1D ^1^H NMR spectra of p47A1 and p47A1_core_ in the absence and presence of 2 molar equivalence of p47_171–266_. Emerging resonances arising from complex formation are indicated by black arrows. (C) Overlay of 2D ^1^H–^15^N HSQC spectra of [U-^15^N]-labelled p47_101–266_ in the free state (dark grey) and in the presence of p47A1 (blue) or p47A1_core_ (magenta). (D) Relative emission intensity (*I*_bound_/*I*_free_) of the AnIm probe at 361 nm upon the addition of p47_101-266_ across different labeling positions within p47A1. Error bars represent standard deviations derived from triplicate measurements (n = 3, \**p* < 0.05, \*\**p* < 0.01). (E) Raw ITC thermogram and (F) corresponding integrated binding isotherm for the titration of p47_101-266_ into p47A1mut recorded in 25 mM Tris, 25 mM NaCl and 1 mM EDTA, pH 6.6.

We then sought to identify critical residue-level determinants in the aptamer core that drives protein recognition using site-specific fluorescence spectroscopy. An aniline–imidazole (AnIm) photobase probe (Figure 5A, inset) was individually incorporated at positions 21, 27, 30, and 34 in the aptamer core (Figure 5A, purple spheres; Figure S6) during solid-phase oligonucleotide synthesis.^43^ This probe functions as a sensitive reporter of its immediate local environment through protonation-dependent emission shifts.^43^ In its neutral state, stabilized by a hydrophobic environment or restricted solvent accessibility, the probe exhibits an emission maximum at 361 nm; conversely, exposure to bulk aqueous solvent drives protonation of the imidazole probe, shifting the emission maximum to 407 nm. Upon addition of p47_101-266_, the fluorescent response of the AnIm probe varied depending on its location in the aptamer. Notably, incorporation of the probe at position 30 yielded the largest increase in relative emission intensity at 361 nm upon complex formation with p47 (Figure 5D and S7). The contrast in fluorescent response indicates that position 30 experiences the most profound transition from a solvent-exposed environment into a hydrophobic or shielded environment upon binding to p47 compared to the other evaluated positions, implying that position 30 forms direct interfacial contacts with the protein target.

To further confirm that the region surrounding position C30 was crucial for binding, the 29′-GCGG-32′ sequence was replaced with 29′-AAGA-32′ (Figure 5A, p471A_mut_) to determine whether these substitutions would disrupt binding. Titration of p47_101-266_ into p471A_mut_ yielded substantially reduced heats comparable to the background heats of dilution (Figure 5E), in stark contrast to the titration into p47A1 (Figure 4C). Furthermore, the integrated isotherm lacks a characteristic binding profile, preventing reliable fitting (Figure 5F). These results underscore the critical role of the sequence surrounding C30, suggesting that this localized region either directly engages p47 at the binding interface or structurally stabilizes Stem 2 to maintain a binding-competent conformation.

### The p47A1 aptamer engages p47 in the native p47-p97 complex

A principal cellular role of p47 is regulating intracellular membrane dynamics, such as post-mitotic Golgi reassembly, via assembling into a functional complex with the AAA+ ATPase p97.^26,28,44,45^ We therefore sought to determine whether the p47A1 aptamer can effectively target p47 in the assembled p47-p97 complex. Previous studies have shown that p47 anchors to p97 through multivalent interactions involving its UBX domain, two flanking SHP motifs (SHP_N_ and SHP_C_), and the dynamic UBA-SIM linker (Figure 6A).^39,40,46^ To ensure optimal NMR spectral resolution while retaining all the p47-p97 contacts, subsequent structural characterizations were conducted using a complex comprising p47 and a functional truncated p97_ND1L_ construct containing the NTD and D1 domains (Figure 6A).^47–49^ We first recorded 1D ¹H NMR spectra profiling the imino proton region of the aptamer in the presence of the p47-p97_ND1L_ complex (Figure 6B, red). Addition of the p47-p97 _ND1L_ complex leads to a pronounced attenuation of the sharp imino resonances corresponding to the aptamer’s terminal stem (Figure 6B, black *vs.* red, green arrows), closely mirroring the spectral changes observed upon binding isolated p47 (Figure 6B, blue) and confirming direct interaction between the aptamer and the complex. Notably, the imino peak at 11.5 ppm induced by binding of the isolated p47 were absent in the presence of the p47-p97_ND1L_ complex (Figure 6B, black arrow). This was likely due to severe line broadening of the aptamer signal in the core region resulted from the marked increase in the rotational correlation time upon aptamer recruitment into the ∼440-kDa p47-p97_ND1L_ complex.

**Figure 6.**
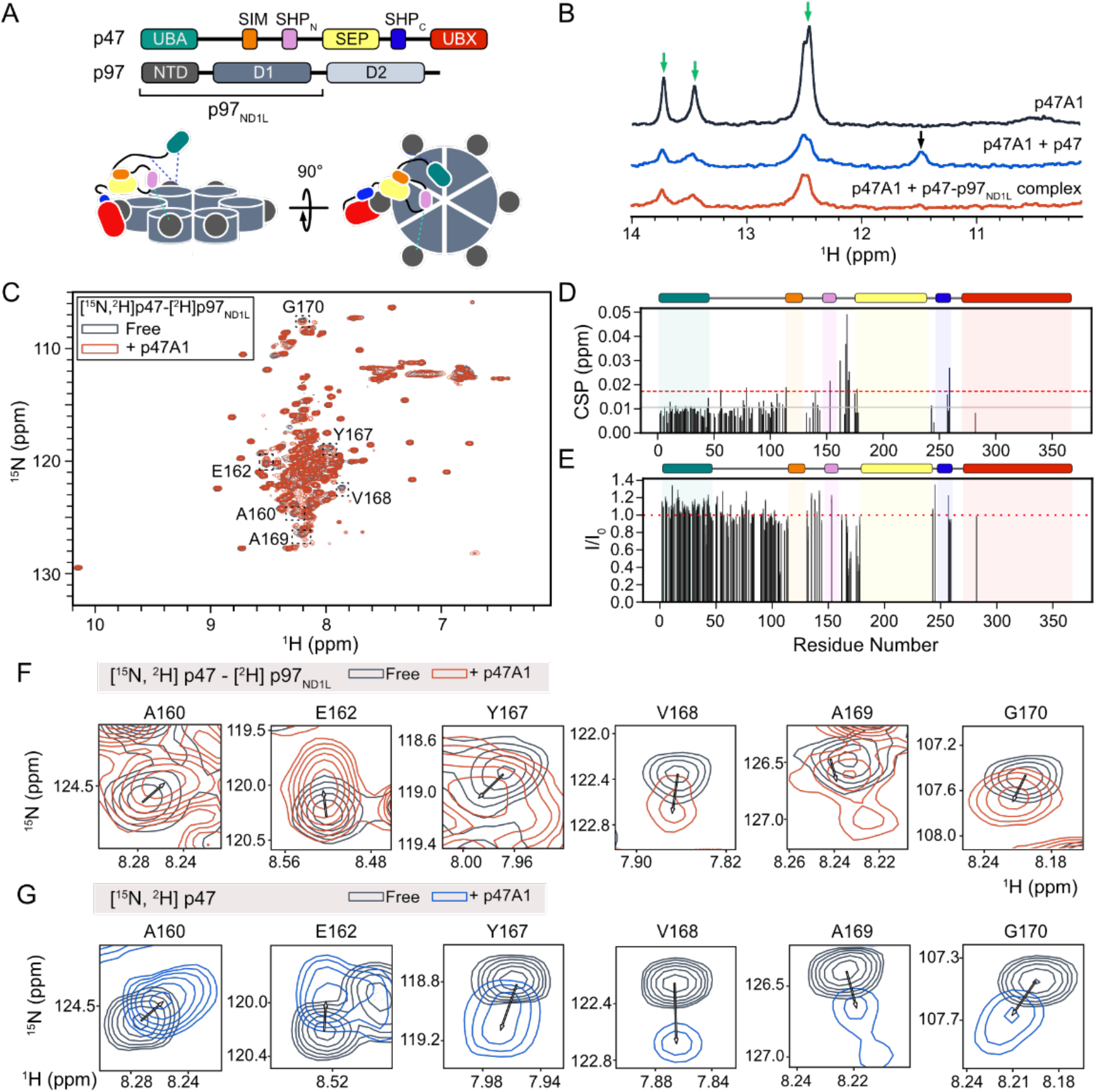
p47A1 interacts with p47 in the p47–p97_ND1L_ complex. (A) Domain architecture of p47, p97 and p97_ND1L_ (top), and a schematic representation of the assembled p47-p97_ND1L_ complex (bottom). Cyan dashed lines represent nucleotide state-dependent interaction between p47 SHP_N_ and p97 NTD,^39^ while blue dashed lines represent transient interactions between p97 NTD and p47 regions.^46^ (B) The imino proton region of 1D ^1^H NMR spectra of p47A1 alone (top), in complex with p47 (middle), and with the pre-assembled p47–p97_ND1L_ complex (bottom). Stoichiometries are maintained at a 1:1 aptamer-to-p47 molar ratio, and a 3:1 p47 monomer-to-p97 hexamer ratio. (C) Overlay of ^1^H-^15^N HSQC spectra of [U-^15^N,^2^H]-p47-[^2^H]-p97_ND1L_ complex in the absence (dark grey) and presence (coral) of the aptamer. Representative residues with significant chemical shift perturbations (CSPs) are highlighted in dashed boxes and labeled. (D) Residue-specific CSPs of [U-^15^N, ^2^H]-p47 in the p47-p97_ND1L_ complex upon aptamer binding plotted from (C). The solid grey and red dashed lines denote the mean and one standard deviation above the mean, respectively. (E) Peak intensity ratios (*I/I₀*) of [U-^15^N, ^2^H]-p47 in the complex between aptamer-bound and free states plotted from (C). The red dotted line indicates no change in peak intensity (*I/I₀* = 1). (F) Closer-up views of selected residues in dashed boxes from (C), showing CSPs between the free (dark grey) and aptamer-bound (coral) states of the p47-p97_ND1L_ complex. (G) Corresponding closer-up spectral regions showing CSPs for isolated [U-^15^N, ^2^H] p47 in the absence (dark grey) and presence (blue) of the aptamer.

We then investigate if the aptamer engages the same structural epitope on p47 in the p47-p97_ND1L_ complex as it does with the isolated p47. We recorded 2D ^1^H-^15^N HSQC spectra of [U-^15^N, ^2^H]-labeled p47 pre-assembled with [U-^2^H]-label p97_ND1L_ in the absence and presence of the aptamer (Figure 6C). Due to the high molecular weight of the complex, resonances originating from the SEP and UBX domains suffered extensive relaxation broadening, leaving only the highly dynamic UBA domain and flexible linkers visible (Figure 6C-E). Upon the addition of the aptamer, chemical shift perturbations were observed for specific residues located on the SIM-SEP linker, including A160, E162, Y167, V168, A169 and G170 (Figure 6D&F). Importantly, the chemical shift perturbation trajectories of these individual residues closely resemble those observed during titrations with isolated p47 (Figure 6F *v.s*. 6G), strongly supporting that the binding epitope of the aptamer on p47 is preserved within the p47-p97_ND1L_ complex. Concurrently, we observed intensity gains for the UBA domain and the UBA-SEP linker upon aptamer binding to the complex (Figure 6E), which mirrors the behavior observed with the isolated p47 (Figure 3D). This suggests that competitive aptamer docking to the SEP domain similarly displaces the intramolecular interactions within p47 and literates the UBA domain, a binding mode preserved in the context of the p47-p97 complex.

To determine whether aptamer binding perturbs the enzymatic function of the complex, we evaluated the steady-state ATP hydrolysis rate (Figure S8). The basal ATPase activity of the p47-p97 complex is largely unaffected upon titration with equal molar or excess quantities of the p47A1 aptamer (Figure S8). This functional preservation is highly consistent with our structural model. Because p47A1 selectively targets the central SIM-SEP region of p47, it neither impedes p47-p97 assembly, which is predominantly driven by p47 UBX domain, nor perturbs the ATP hydrolysis site of p97 located distantly from the aptamer docking interface. Nonetheless, given that the SIM-SEP region is implicated in substrate engagement,^39,40^ the functional consequences of aptamer binding on downstream substrate unfolding and Golgi reassembly remain to be elucidated via further *in vitro* and *in vivo* studies.

Taken together, these findings demonstrate that the p47A1 aptamer recognizes p47 within the context of the native p47–p97 complex without requiring p47 dissociation, highlighting its potential utility as a chemical biology probe to interrogate and manipulate p47-dependent p97 pathways in a physiologically relevant macromolecular context.

## Conclusion

In this study, we developed and thoroughly characterized p47A1, the first DNA aptamer that targets a regulatory adaptor protein of the p97 AAA+ ATPase. By integrating SELEX, NMR, ITC, site-specific photobase fluorescence, and mutagenesis, we establish a rigorous biophysical framework that elevates protein-aptamer discovery beyond initial screening and characterization. We show that p47A1 recognizes the central SEP domain and adjacent SIM-SEP linker on p47 via specific contacts and maintain robust low-micromolar affinity across a range of ionic conditions. We identified the critical region surrounding C30 on the aptamer that drives complex formation. Mechanistically, we elucidated that the aptamer engagement acts as an allosteric switch, displacing transient intramolecular contacts between the SEP domain and the UBA domain and thereby liberating the UBA domain and potentially remodeling its competence for polyubiquitinated substrate engagement. Crucially, p47A1 successfully engages p47 within the assembled p47-p97 complex while preserving p97’s basal ATPase turnover, validating the vital concept of cofactor-selective targeting over indiscriminate catalytic inhibition. The development of p47A1 provides immediate opportunities for chemical biology applications and therapeutic innovation, such as in-cell fluorescence imaging tools to monitor real-time p47 localization and Golgi dynamics, and aptamer-based proteolysis-targeting chimeras to induce selective destruction of p47 in cancer cells. More broadly, this study showcases a generalizable strategy for deploying oligonucleotides to dissect and modulate the expansive repertoire of multi-domain adaptors governing the functions of AAA+ ATPases and related macromolecular machines.

## Supporting Information

The authors have cited additional references within the Supporting Information.

## Data availability

All processed data are available in the article and supporting information. NMR raw data are available upon request.

## Acknowledgements

Funding for this work was provided by the Natural Sciences and Engineering Research Council of Canada (NSERC) Discovery Grants RGPIN-2020-00527 to R.H., RGPIN 2020-05043 to D.O., RGPIN-2026-04355 to J.L., and RGPIN-2025-04087 to R.M.. We acknowledge Cancer Research Society Operating Grant to R.H. for providing salary support for L.F.. We also acknowledge the Canada Research Chairs program’s support to J.L.. P.E.K acknowledges support for an NSERC Canada Graduate Scholarship (CGS M). We acknowledge the support from CFI-JEFL grants (43757 and 45118) to R.H, D.O. and R.M. that provided the equipment access to this study. NMR data were recorded at the University of Guelph NMR Center. We thank Dr.Sameer Al-Abdul-Wahid (University of Guelph) for support with NMR measurements. We thank Nicholas Horton for his help and suggestions with the purification of the aptamer.

## Conflicts of Interests

The authors declare no conflict of interest.

## Notes

### Competing Interest Statement

The authors have declared no competing interest.

