## Supplementary Material for "Structural and Biophysical Basis of the First DNA Aptamer Targeting the Dynamic p47 Adaptor of the p97 AAA+ Machine"

**Protein sequences**

**p47$_{WT}$**

40709.98 Da

Predicted pI: 5.04

GMAEERQDALREFVAVTGTEEDRARFFLESAGWDLQIALASFYEDGGDEDIVTISQATPSSVSRGTAPSD
NRVTSFRDLIHDQDEEEEEEEEGQRFYAGGSERSGQQIVGPPRKKSPNELVDDLFKGAKEHGAVAVERVT
KSPGETSKPRPFAGGGYRLGAAPEEESAYVAGERRRHSGQDVHVVLKLWKTGFSLDNGDLRSYQDPSN
AQFLESIRRGEVPAELRRLAHGGQVNLDMEDHRDEDFVKPKGAFKAFTGEGQKLGSTAPQVLNTSSPAQ
QAENEAKASSSILINEAEPTTNIQIRLADGGRLVQKFNHSHRISDIRLFIVDARPAMAATSFVLMTTFPNKEL
ADENQTLKEANLLNAVIVQRLT

**p47$_{101\text{-}266}$**

18147.20 Da

Predicted pI: 8.87

GRSGQQIVGPPRKKSPNELVDDLFKGAKEHGAVAVERVTKSPGETSKPRPFAGGGYRLGAAPEEESAY
VAGERRRHSGQDVHVVLKLWKTGFSLDNGDLRSYQDPSNAQFLESIRRGEVPAELRRLAHGGQVNLDM
EDHRDEDFVKPKGAFKAFTGEGQKLGSTAPQ

**p47$_{171\text{-}266}$**

10832.02 Da

Predicted pI: 7.19

ERRRHSGQDVHVVLKLWKTGFSLDNGDLRSYQDPSNAQFLESIRRGEVPAELRRLAHGGQVNLDMEDH
RDEDFVKPKGAFKAFTGEGQKLGSTAPQ

**p97$_{ND1L}$**

53437.26 Da

Predicted pI: 5.18

GMASGADSKGDDLSTAILKQKNRPNRLIVDEAINEDNSVVSLSQPKMDELQLFRGDTVLLKGKKRREAVCI
VLSDDTCSDEKIRMNRVVRNNLRVRLGDVISIQPCPDVKYGKRIHVLPIDDTVEGITGNLFEVYLKPYFLEA
YRPIRKGDIFLVRGGMRAVEFKVVETDPSPYCIVAPDTVIHCEGEPIKREDEEESLNEVGYDDIGGCRKQL
AQIKEMVELPLRHPALFKAIGVKPPRGILLYGPPGTGKTLIARAVANETGAFFFLINGPEIMSKLAGESESNL
RKAFEEAEKNAPAIIFIDELDAIAPKREKTHGEVERRIVSQLLTLMDGLKQRAHVIVMAATNRPNSIDPALRR
FGRFDREVDIGIPDATGRLEILQIHTKNMKLADDVDLEQVANETHGHVGADLAALCSEAALQAIRKKMDLID
LEDETIDAEVMNSLAVTMDDFRWALSQSNPSALRETVVEVPQVTWEDIG

### Supplementary Tables

**Table S1.** Selection conditions used for each round.

| Round No. | Library (pmol) | Beads volume (μL) | Buffer wash (times) |
|---|---|---|---|
| 1 | 100 | 20 | 3 |
| 2 | 50 | 20 | 3 |
| 3 | 50 | 20 | 3 |
| 4 | 50 | 20 | 3 |
| 5 | 50 | 20 | 3 |
| 6 | 50 | 20 | 3 |
| 7 | 50 | 20 | 3 |
| 8 | 50 | 20 | 3 |
| 9 | 50 | 20 | 5 |
| 10 | 50 | 20 | 5 |

**Table S2.** Fitted parameters for DLS autocorrelation functions.

| Parameters | p47A1 pre-annealing no $Mg^{2+}$ | p47A1 post-annealing no $Mg^{2+}$ | p47A1core post-annealing no $Mg^{2+}$ | p47A1 post-annealing 5 mM $Mg^{2+}$ |
|---|---|---|---|---|
| Diffusion coefficient ($m^2/s$) | $4.28e^{-12}$ | $1.14e^{-10}$ | $1.29e^{-10}$ | $1.15e^{-10}$ |
| Diffusion coefficient error ($m^2/s$) | $1.29e^{-13}$ | $3.22e^{-13}$ | $4.14e^{-13}$ | $3.07e^{-13}$ |
| $\mu^2$ | $3.26e^{6}$ | $5.58e^{8}$ | $5.12e^{8}$ | $5.49e^{8}$ |
| $\Gamma$ ($s^{-1}$) | $1.65e^{3}$ | $4.39e^{4}$ | $4.98e^{4}$ | $4.43e^{4}$ |
| PD | 1.191 | 0.290 | 0.207 | 0.280 |
| $R_h$ (m) | $5.74e^{-8}$ | $2.16e^{-9}$ | $1.91e^{-9}$ | $2.14e^{-9}$ |
| $R_h$ error (m) | $1.74e^{-9}$ | $6.13e^{-12}$ | $6.13e^{-12}$ | $5.73e^{-12}$ |

**Table S3.** Thermodynamic parameters of binding of the p47A1 aptamer or its truncated variant p47A1$_{core}$ to the three p47 variants determined by ITC. Measurements were performed in Buffer A (25 mM Bis-Tris, 25 mM NaCl, 1 mM EDTA, pH 6.6), Buffer B (20 mM MES, 150 mM NaCl, 5 mM $MgCl_2$, 10 mM KCl, pH 6.2), and Buffer C (25 mM Bis-Tris, 25 mM NaCl, 1 mM EDTA, 5 mM $MgCl_2$, pH 6.6).

| **p47 variant** | **Aptamer variant** | **Buffer Condition** | **$K_d$ (μM)** | **$\Delta G$ (kcal·mol$^{-1}$)** | **$\Delta H$ (kcal·mol$^{-1}$)** | **$-T\Delta S$ (kcal·mol$^{-1}$)** | ***N*** |
|---|---|---|---|---|---|---|---|
| p47$_{WT}$ | p47A1 | Buffer A | 9.8 ± 4.8 | -6.8 | -4.4 ± 2.4 | -2.7 | 0.50 ± 0.16 |
| p47$_{101\text{-}266}$ | p47A1 | Buffer A | 1.6 ± 0.7 | -7.9 | -4.5 ± 0.3 | -3.4 | 0.97 ± 0.05 |
| p47$_{171\text{-}266}$ | p47A1 | Buffer A | 4.1 ± 1.5 | -7.4 | -4.5 ± 0.4 | -2.8 | 1.2 ± 0.1 |
| p47$_{WT}$ | p47A1 | Buffer B | 4.1 ± 1.7 | -7.4 | -4.7 ± 0.6 | -2.6 | 1.1 ± 0.1 |
| p47$_{101\text{-}266}$ | p47A1 | Buffer B | 12 ± 5 | -6.7 | -5.6 ± 2.0 | -1.2 | 0.77 ± 0.20 |
| p47$_{171\text{-}266}$ | p47A1 | Buffer B | 3.9 ± 1.3 | -7.4 | -8.7 ± 1.0 | 1.3 | 0.74 ± 0.06 |
| p47$_{WT}$ | p47A1 | Buffer C | 5.9 ± 2.3 | -7.1 | -5.9 ± 1.1 | -1.3 | 0.89 ± 0.12 |
| p47$_{101\text{-}266}$ | p47A1 | Buffer C | 2.7 ± 1.0 | -7.6 | -6.4 ± 0.7 | -1.2 | 0.74 ± 0.06 |
| p47$_{171\text{-}266}$ | p47A1 | Buffer C | 1.9 ± 0.8 | -7.8 | -10 ± 1 | 2.4 | 0.79 ± 0.06 |
| p47$_{101\text{-}266}$ | p47A1$_{core}$ | Buffer A | 2.6 ± 0.8 | -7.6 | -15 ± 1 | 7.2 | 1.5 ± 0.1 |

**Table S4.** Experimental conditions for the ITC titrations. Aptamers were placed in the sample cell, and p47 variants were titrated from the syringe at the concentrations listed. All titrations were performed at 25 °C with 20 injections of the indicated volume, spaced 300 s apart.

| p47 variant | Aptamer variant | Buffer Condition | Aptamer [Cell] (μM) | p47 variants [Syringe] (μM) | Injection volume (μL) |
|---|---|---|---|---|---|
| $p47_{WT}$ | p47A1 | Buffer A | 25 | 235 | 2.5 |
| $p47_{101-266}$ | p47A1 | Buffer A | 25 | 250 | 2 |
| $p47_{171-266}$ | p47A1 | Buffer A | 20 | 250 | 2.5 |
| $p47_{WT}$ | p47A1 | Buffer B | 20 | 200 | 2.5 |
| $p47_{101-266}$ | p47A1 | Buffer B | 25 | 250 | 2.5 |
| $p47_{171-266}$ | p47A1 | Buffer B | 25 | 250 | 2.5 |
| $p47_{WT}$ | p47A1 | Buffer C | 20 | 250 | 2 |
| $p47_{101-266}$ | p47A1 | Buffer C | 25 | 250 | 2 |
| $p47_{171-266}$ | p47A1 | Buffer C | 25 | 250 | 2 |
| $p47_{101-266}$ | $p47A1_{core}$ | Buffer A | 25 | 230 | 2.5 |

**Table S5.** DNA sequences used in the selection.

| DNA Name | Sequences (From 5′ to 3′) |
|---|---|
| Library | GGAGGCTCTCGGGACGAC**$N_{30}$**GTCGTCCCGACTCTATGATGACTGT |
| Forward Primer | GGAGGCTCTCGGGACGAC |
| Biotin-Reverse | /5Biosg/ACAGTCATCATAGAGTCGGGACG |
| P5-503 | AATGATACGGCGACCACCGAGATCTACACTATCCTCTACACTCTTTCCCTACA CGACGCTCTTCCGATCTACAGTCATCATAGAGTCGGGACG |
| P7-705 | CAAGCAGAAGACGGCATACGAGATAGGAGTCCGTGACTGGAGTTCAGACGT GTGCTCTTCCGATCTGGAGGCTCTCGGGACGAC |

**Table S6.** Experimental buffer compositions.

| Buffer Number | Experimental | Buffer Components |
|---|---|---|
| 1 | NMR/ITC/DLS | 25 mM Bis-Tris (pH 6.6), 25 mM NaCl, 1 mM EDTA, 5% $D_2O$ |
| 2 | NMR/ITC/DLS | 25 mM Bis-Tris (pH 6.6), 25 mM NaCl, 1 mM EDTA, 5 mM $MgCl_2$, 10 mM KCl, 5% $D_2O$ |
| 3 | ITC | 20 mM MES (pH 6.2), 150 mM NaCl, 5 mM $MgCl_2$, 10 mM KCl |
| 4 | CD/$T_m$ | 10 mM $Na_2HPO_4$, 1.8 mM $KH_2PO_4$ (pH 6.6), 25 mM NaCl |
| 5 | CD/$T_m$ | 10 mM $Na_2HPO_4$, 1.8 mM $KH_2PO_4$ (pH 6.6), 25 mM NaCl, 10 mM $MgCl_2$ |
| 6 | Fluorescence | 50 mM Tris-HCl (pH 7.7), 25 mM NaCl |

**Supplementary Figures**

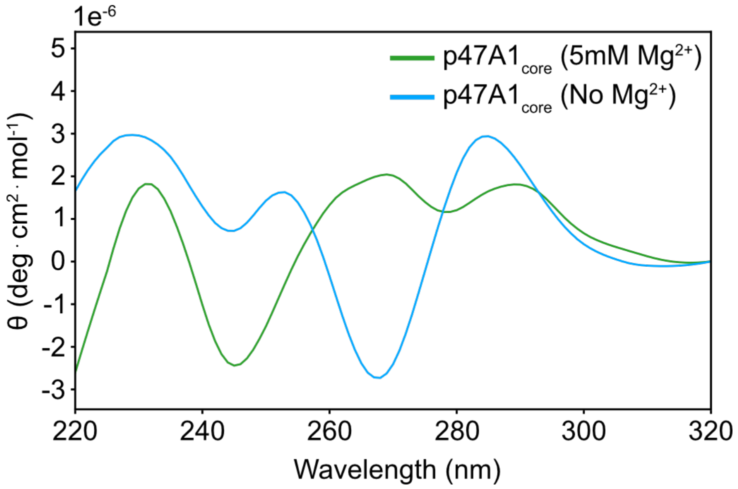


**Figure S1.** CD spectra of p47A1$_{core}$ in the absence (cyan) and presence (green) of 5 mM $MgCl_2$. Measurements were performed in a buffer containing 10 mM $Na_2HPO_4$, 1.8 mM $KH_2PO_4$ (pH 6.6), 25 mM NaCl, and 5 mM $MgCl_2$ (where indicated).

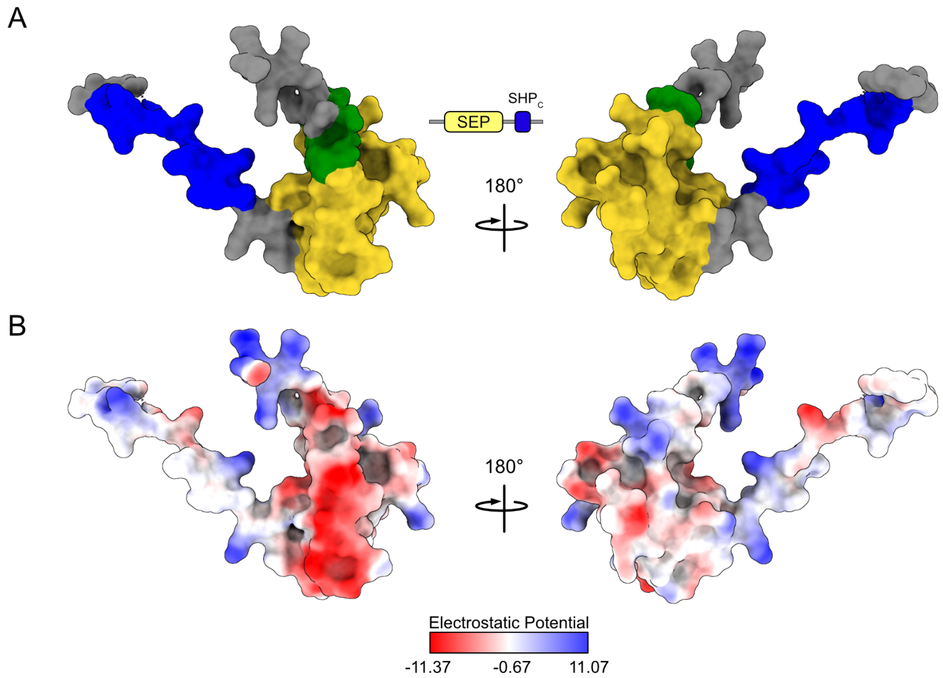


**Figure S2.** Binding interface of the p47A1 aptamer on $p47_{171-266}$. (A) Surface representation of $p47_{171-266}$ with residues involved in aptamer binding highlighted in green. (B) Electrostatic surface of $p47_{171-266}$ illustrates the binding interface consists of both acidic and hydrophobic residues.

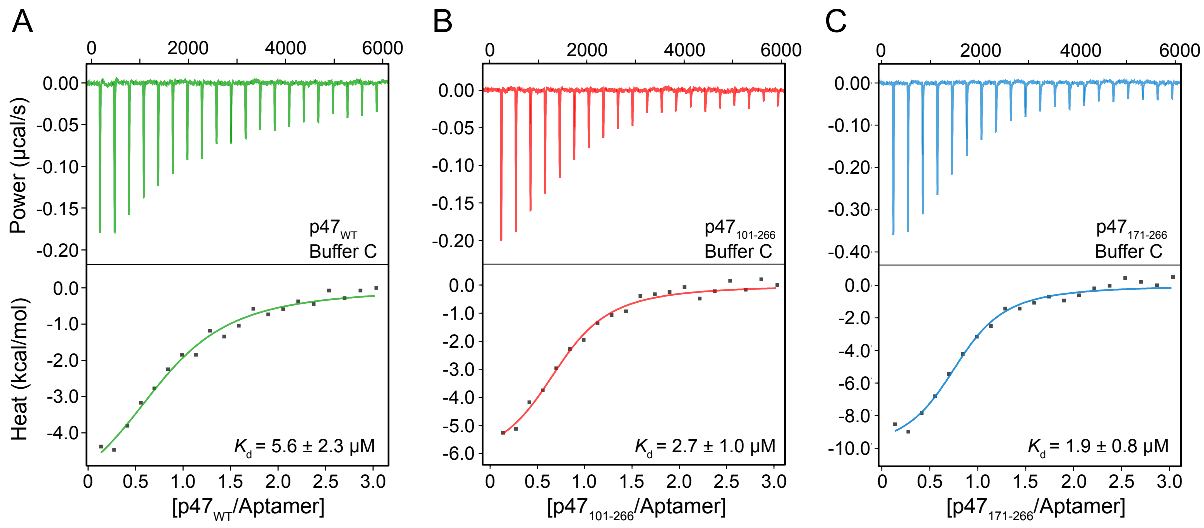


**Figure S3.** Representative raw ITC thermograms (top) and corresponding integrated binding isotherms (bottom) for titrations of $p47_{WT}$ (A), $p47_{101–266}$ (B), and $p47_{171-266}$ (C) into the p47A1 aptamer in Buffer C (25 mM Tris, 25 mM NaCl, 1 mM EDTA, 5 mM $MgCl_2$, pH 6.6). Solid lines in the binding isotherms represent the best fit to a two-state N-equivalent site binding model, with the fitted parameters summarized in Table S3.

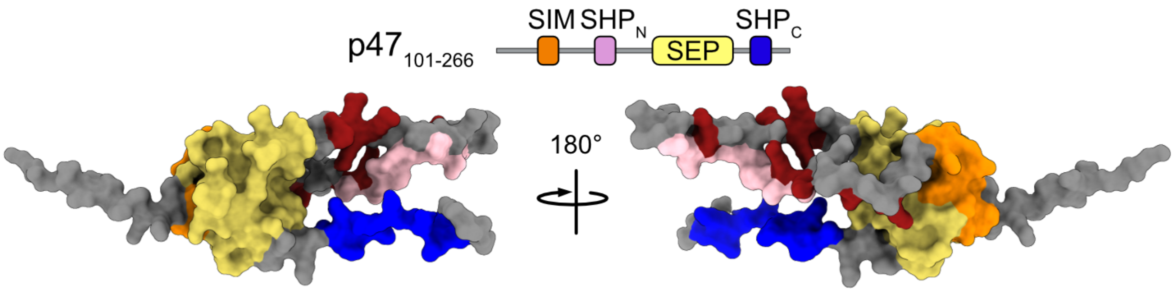


**Figure S4.** Surface representation of $p47_{101–266}$, with positively charged residues highlighted in red.

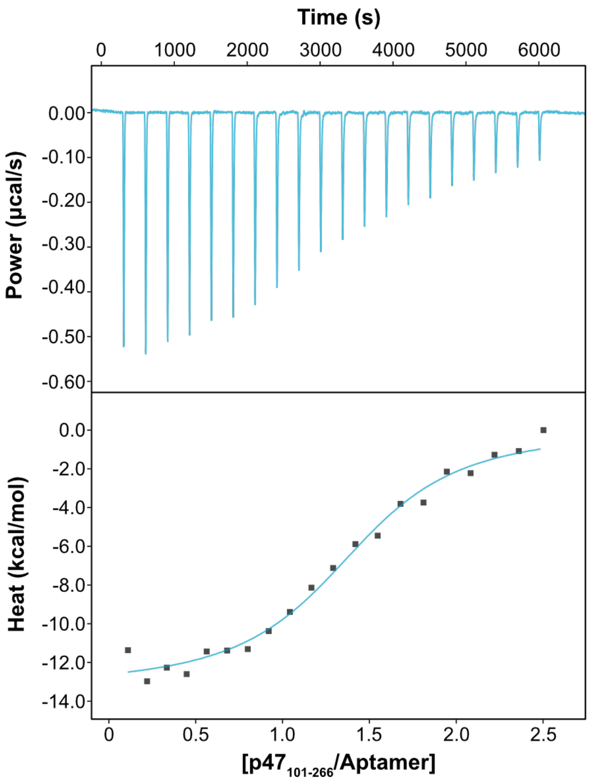


**Figure S5.** Representative raw ITC thermograms (top) and corresponding integrated binding isotherms (bottom) for titrations of $p47_{101–266}$ into the $p47A1_{core}$ aptamer in a buffer containing 25 mM Tris, 25 mM NaCl, 1 mM EDTA, pH 6.6. Solid lines in the binding isotherms represent the best fit to a two-state N-equivalent site binding model, with the fitted parameters summarized in Table S3.

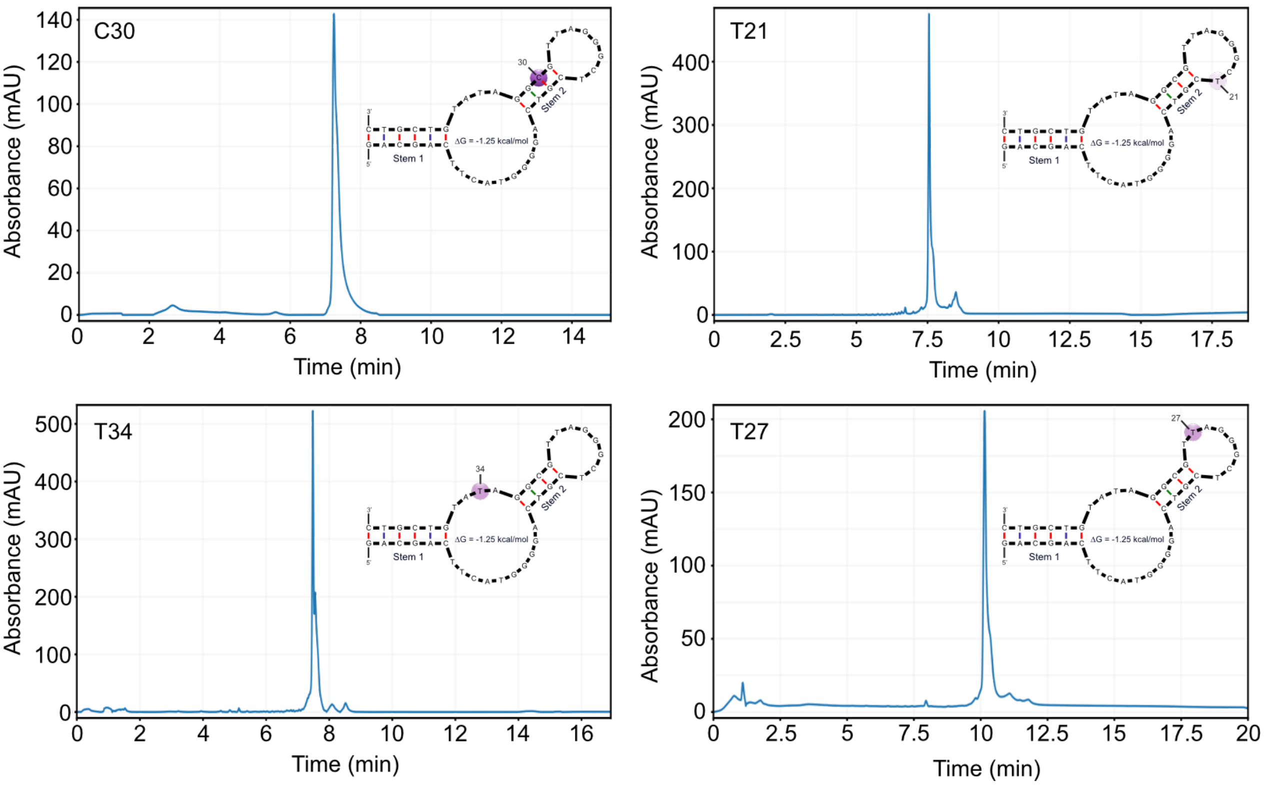


**Figure S6.** SAX HPLC profiles of aptamer preparations with AnIm incorporated at positions 21, 27, 30, and 34. The column was eluted using a linear gradient of 0–100% buffer B over 15 min for positions 21, 30, and 34, and over 20 min for position 27 (buffer A: 50 mM Tris HCl, pH 7.61, 10% acetonitrile; buffer B: 50 mM Tris HCl, pH 7.61, 10% acetonitrile, 1 M NaCl) at 55 °C and a flow rate of 1 mL/min. Monitoring was performed at 260 nm.

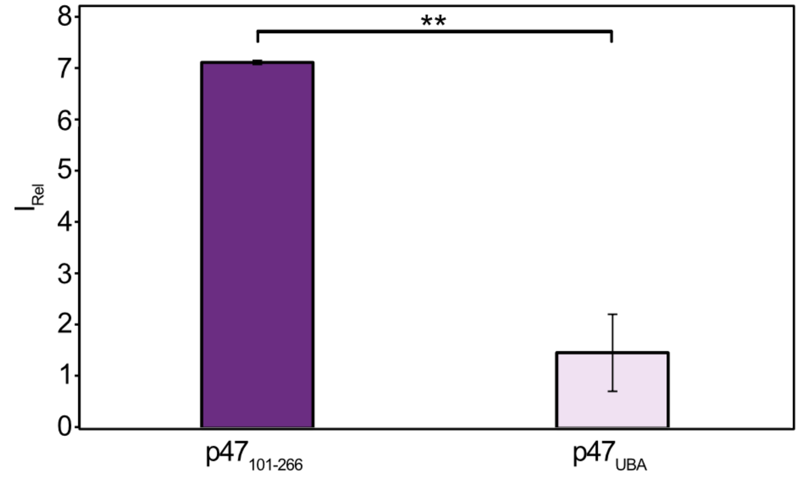


**Figure S7.** Relative emission intensity ($I_{Rel} = I_{bound}/I_{free}$) of the AnIm probe at 361 nm upon addition of $p47_{101-266}$ or $p47_{UBA}$. Error bars represent standard deviations derived from triplicate measurements ($n = 3$, $^{**}p < 0.01$).

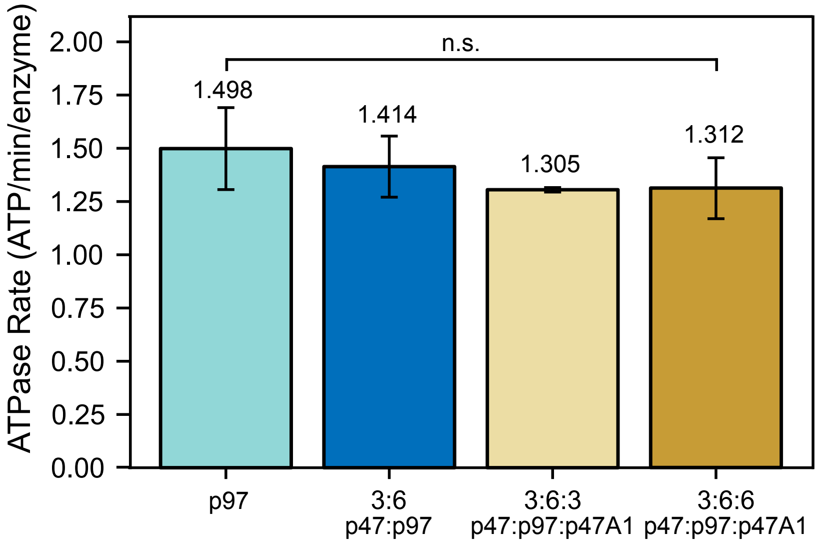


**Figure S8.** ATP hydrolysis rates of p97 alone (light blue), the p47-p97 complex (blue), and the p47-p97 complex in the presence of an equimolar equivalent (light yellow) or 2-fold molar access (yellow) of the p47A1 aptamer with respect to p47. No significant difference (n.s.) was observed between the four samples.

### Experimental Section

#### Materials

All DNA oligonucleotides (ONs) used for selection were purchased from Integrated DNA Technologies (Coralville, IA, USA), and their sequences are listed in Table S5. 2-(*N*-morpholino)ethanesulfonic acid (MES), NaCl, $MgCl_2$, and KCl were obtained from BioShop Canada. SsoFast EvaGreen Supermix was purchased from Bio-Rad. Taq DNA polymerase, dNTPs, and Taq reaction buffer were obtained from New England Biolabs. NHS-activated agarose resin and streptavidin-coated agarose resin (1–3 mg biotinylated BSA per mL resin) were purchased from Thermo Scientific.

#### Protein Expression and Purification

Full-length and truncated *Mus musculus* p47 constructs (*i.e.* $p47_{101-266}$, $p47_{171-266}$) were expressed from a pET29 vector. Full-length *Mus musculus* p97 and $p97_{ND1L}$ were expressed from a pET28b vector. All constructs contained an *N*-terminal $His_6$ tag followed by a tobacco etch virus (TEV) protease cleavage site. All proteins were expressed in *Escherichia coli* BL21(DE3) cells and purified following previously established protocols[1–3]. [U-$^{15}$N] p47 constructs were expressed in M9 minimal medium supplemented with $^{15}NH_4Cl$ (299251; Milli-pore Sigma) as the sole nitrogen source, while uniformly [$^2$H] $p97_{ND1L}$ and [U-$^{15}$N, $^2$H] p47 were produced in $D_2O$-based M9 media with [$^2$H,$^{12}$C]-glucose as the carbon source, as previously described[1,3]. Purified proteins were exchanged into the appropriate experimental buffers (Table S6) using Amicon Ultra centrifugal filters (Millipore), flash-frozen in liquid nitrogen, and stored at −80 °C. Protein purity (>95%) was confirmed by SDS-PAGE, and protein concentrations were determined from the absorbance at 280 nm using extinction coefficients calculated with ExPASy ProtParam.

#### Target-Immobilized SELEX

NHS-activated agarose beads (125 µL) were washed sequentially with Milli-Q water (2 mL) and 1× PBS (2 mL; 10 mM phosphate, 150 mM NaCl) before incubation with 300 µL of 28 µM full-length p47 ($p47_{WT}$) for 1 h at room temperature with end-over-end mixing. Residual NHS groups were quenched with 500 µL of 1 M ethanolamine (pH 7.4) for 15 min, after which the beads were washed extensively with 1× PBS, resuspended in 200 µL of 1× PBS, and stored at 4 °C until use.

A 73-nt ssDNA library containing a 30-nt randomized region was used for SELEX. Before each round, 100 pmol of the DNA library was annealed by heating to 95 °C for 2 min followed by slow cooling to room temperature. The annealed library was incubated with 20 µL of p47-conjugated agarose beads in selection buffer (20 mM MES, pH 6.2, 150 mM NaCl, 5 mM $MgCl_2$, 10 mM KCl, and 0.005% (v/v) Tween-20) for 30 min at room temperature with end-over-end mixing. Following incubation, the beads were washed three times with 200 µL of selection buffer, and bound DNA was eluted with 120 µL of elution buffer (10 mM EDTA, 10 mM Tris-HCl, pH 8.4). The recovered DNA was amplified by PCR using an initial denaturation at 95 °C for 5 min, followed by 13–15 cycles of 95 °C for 15 s, 55 °C for 30 s, and 68 °C for 30 s, with a final extension at 68 °C for 3 min.

PCR products were converted to ssDNA by immobilization on a column containing 150 µL of agarose beads and 50 µL of streptavidin-coated agarose resin. Following washing with ten 500 µL aliquots of 1× PBS, strand separation was achieved by incubation with 400 µL of 0.2 M NaOH for 5 min, followed by an additional 200 µL of 0.2 M NaOH. The combined eluates were neutralized with 600 µL of 0.2 M HCl, concentrated using a 10-kDa molecular weight cut-off centrifugal filter, washed three times with Milli-Q water, and concentrated to a final volume of 60 µL. DNA concentration was determined by NanoDrop spectrophotometry, and the recovered ssDNA was used as the input library for the subsequent round of selection. Selection conditions for each round are summarized in Table S1.

**DNA Aptamer Synthesis and Purification**

DNA Oligonucleotides (ONs) were synthesized on an ABI 394 DNA synthesizer using standard phosphoramidite chemistry. dA, dG, dC, and dT phosphoramidites and all synthesis reagents were purchased from ChemGenes (Wilmington, MA) and used according to the manufacturer's protocols. Following synthesis, ONs were cleaved from the solid support and deprotected with a 4:1 (v/v) mixture of 25% ammonium hydroxide and ethanol (0.5–1.0 mL) for 17 h at 55 °C. Samples were partially concentrated under vacuum using a Savant SC110A SpeedVac to remove residual volatile base. Deprotected ONs were purified by strong anion-exchange (SAX) HPLC on a DNAPac™ PA200 column using a Vanquish™ analytical purification HPLC system with a linear gradient of 0–100% elution buffer (1 M NaCl in 10% acetonitrile, 50 mM Tris, pH 7.61). Purified ONs were desalted using Sep-Pak C18 Classic cartridges (Waters), which were conditioned sequentially with acetonitrile, 50% acetonitrile/water, and 100 mM sodium acetate (pH 7.0). Samples were diluted to ≤2% acetonitrile prior to loading, washed with Milli-Q water, eluted with 50% acetonitrile/water, and concentrated under vacuum. ON concentrations were determined from absorbance at 260 nm using an Agilent BioTek Epoch microplate spectrophotometer. Each sample was measured in triplicate, blank corrected, and quantified using the Beer–Lambert equation with molar extinction coefficients calculated using the OligoAnalyzer™ Tool (Integrated DNA Technologies). Prior to use, aptamers were diluted to 100–150 µM in the appropriate annealing buffer (Table S6), heated to 95 °C for 10 min, and allowed to cool gradually to room temperature to promote folding. Annealed aptamers were stored at 4 °C for short-term use or at −20 °C for long-term storage.

**Dynamic Light Scattering**

Dynamic light scattering (DLS) experiments were carried out on a DynaPro DLS Plate Reader III (Wyatt Technology). Prior to DLS measurements, samples were centrifuged at 20,000 × g for 12 min at 25 °C to remove aggregates and particulates. The supernatants were then transferred to a 384-well plate, and the plate was centrifuged at 3,000 × g for 2 min at 4 °C to remove air bubbles. Measurements were performed at 25 °C with 20 acquisitions collected per well and averaged for analysis. The autocorrelation functions were subjected to cumulant fitting to obtain the translational diffusion coefficient (D) as previously described.[4,5] The fitted diffusion coefficient, D, was converted to the hydrodynamic radius, $R_\mathrm{h}$, using the Stokes-Einstein equation

$$R_\mathrm{h} = \frac{k_\mathrm{B}T}{6\pi\eta D}$$

where $k_\mathrm{B}$ is the Boltzmann constant, $T$ is the temperature in Kelvin, and η is the viscosity.[6] Polydispersity (PD) was calculated as $PD = \frac{\mu_2}{\Gamma^2}$, where $\Gamma = Dq^2$ is the mean decay rate and $\mu_2$ is the second cumulant obtained from the fit, following the methods of cumulants.[5] Representative fitting parameters are summarized in Table S2.

#### Circular Dichroism Spectroscopy

Circular dichroism (CD) spectra were acquired using a Jasco J-815 spectropolarimeter equipped with a Julabo F25 circulating water bath for temperature control. Aptamer samples (10 μM) were prepared in buffers 4 and 5 (Table S6) and equilibrated at 25 °C for 5–10 min prior to data collection. Spectra were recorded from 320 to 220 nm in a 1-cm path length quartz cuvette (Starna) using a scan rate of 200 nm • $\mathrm{min}^{-1}$, a 1-nm bandwidth, and a 0.2-nm data pitch. Each spectrum represents the average of five consecutive scans. Baseline corrections were performed by subtracting corresponding buffer spectra collected under identical experimental parameters. Raw ellipticity was converted to molar ellipticity (deg • $\mathrm{cm}^2$ • $\mathrm{mol}^{-1}$) and the spectra were processed using Spectra Manager software suite (JASCO). For visualization, spectra were smoothed with a digital Savitzky–Golay filter prior to plotting.

#### UV-Vis Monitored Thermal Denaturation

Thermal denaturation experiments were performed using a Cary 3500 Multicell UV–Vis Spectrophotometer (Agilent Technologies) equipped with a Peltier temperature controller. All measurements were conducted in 10-mm path length quartz cuvettes with buffer baseline correction. DNA samples were diluted to a final concentration of 4 μM in buffers 4 and 5 (Table S6). Thermal melting profiles were monitored by tracking the absorbance at 260 nm during five alternating heating and cooling cycles between 25 and 90 °C at a ramp rate of 0.5 °C • $\mathrm{min}^{-1}$. Melting temperatures ($T_m$) were determined from the maxima of the first derivative of the hyperchromic transition curves using Cary UV Workstation Software (Agilent Technologies) and are reported as the average of the five thermal cycles.

### NMR Experiments

All NMR experiments were acquired on a 600 MHz Bruker Avance III NMR spectrometer equipped with a 5 mm TCI cryoprobe. One-dimensional $^1$H NMR spectra were recorded using a standard 1D $^1$H ZGESP (excitation sculpting) pulse sequence to achieve water suppression. For 1D $^1$H spectra, aptamer samples (50–100 μM) were prepared in buffers 1 and 2 (Table S6) supplemented with 5% $D_2O$ for field locking. All 1D spectra were recorded at 5 °C with 512 transients and were subsequently processed and analyzed using TopSpin 4.4.1 (Bruker).

Two-dimensional $^1$H–$^{15}$N HSQC spectra were acquired at 25 °C using an interscan delay of 1.5 s. Spectra were recorded with acquisition times of 39 ms ($^{15}$N) and 64 ms ($^1$H), and 70 and 640 complex points in the indirect ($^{15}$N) and direct ($^1$H) dimensions, respectively. The $^1$H and $^{15}$N carrier frequencies were centred on the water resonance (4.67 ppm) and 119 ppm, respectively. Spectra were processed using NMRPipe[7] and analysed using NMRFAM-SPARKY[8].

To characterize the interaction between p47 and the aptamer, $^1$H–$^{15}$N HSQC experiments were performed on $^{15}$N-labelled p47 constructs, including $p47_{WT}$, $p47_{101–266,}$ and $p47_{171–266}$ at a concentration of 60 μM, in the absence and presence of the full-length or truncated p47A1 aptamer at 1:1 molar ratio in a buffer that contains 25 mM Bis-Tris (pH 6.6), 25 mM NaCl, and 1 mM EDTA, and 5 % $D_2O$.

To characterize the p47/p97 complex, $^1$H–$^{15}$N HSQC experiments were performed using [U-$^{15}$N, $^2$H] $p47_{WT}$ in complex with [$^2$H]-labelled $p97_{ND1L}$. $^{15}$N-labelled $p47_{WT}$ (50 μM) was incubated with 100 μM [$^2$H]-labelled $p97_{ND1L}$ (monomer concentration) under the same buffer conditions described above. Following characterization of the p47/p97 complex, p47A1 aptamer was added to the preformed complex at final concentrations of 50 and 100 μM. Subsequent $^1$H–$^{15}$N HSQC spectra were acquired at 298 K to monitor changes in p47 resonance chemical shifts and intensities upon aptamer addition.

Average chemical shift perturbations (CSPs) were calculated as

$$\mathrm{CSP_{av}} = \{\Delta\delta_{\mathrm{H}}{}^2 + (0.14 \times \Delta\delta_{\mathrm{N}})^2\}^{0.5} \qquad (1)$$

where $\Delta\delta_H$ and $\Delta\delta_N$ are the chemical shift differences at the $^1$H and $^{15}$N dimensions, respectively.[9] Binding-induced peak intensity changes were determined by taking the ratio between the individual peak heights of the resonances in the bound state ($I$) against those of the corresponding resonances in the free state ($I_0$). Uncertainty of $I/I_0$ was estimated by propagating the RMS baseline noise of each spectrum through the intensity ratio, according to

$$\sigma_{\frac{I}{I_0}} = \left(\frac{I}{I_0}\right)\sqrt{\left(\frac{\sigma_{free}}{I_0}\right)^2 + \left(\frac{\sigma_{bound}}{I}\right)^2} \qquad (2)$$

where $\sigma_{free}$ and $\sigma_{bound}$ are the RMS baseline noise levels of the free and bound spectra, respectively, taken as an estimate of the standard deviation of individual peak intensities in each spectrum.

**Isothermal titration calorimetry (ITC)**

Isothermal titration calorimetry (ITC) experiments were performed on an Affinity ITC calorimeter (TA Instruments). Prior to titration, protein and aptamer samples were buffer exchanged into the same designated buffer (Table S6) using Amicon centrifugal concentrators to minimize buffer mismatch. Samples were then centrifuged at 20,000 × g for 5 min to pellet any aggregates before loading into the calorimeter. The sample cell (effective volume 185 µL) was loaded with aptamer (p47A1 or p47A1$_{core}$, 20–25 µM), and the syringe was filled with p47 variants (p47$_{WT}$, p47$_{101–266}$ or p47$_{171–266}$, 200–250 µM). Exact concentrations for each titration are listed in Table S4. All measurements were carried out at 25 °C with a stirring rate of 125 rpm. Each titration consisted of 20 injections of 2.0 or 2.5 µL (Table S4) spaced 300 s apart to allow thermal equilibration. Thermograms were baseline corrected, integrated and corrected for heats of dilution. Binding isotherms were fitted to an independent (*N*-equivalent) sites model in NanoAnalyze (TA Instruments) to obtain the thermodynamic parameters summarized in Table S3.

**AnIm Fluorescent Probe Experiments**

The AnIm fluorescent photobase-modified oligonucleotides were synthesized via oligonucleotide solid-phase synthesis. Analytical HPLC chromatographs confirming the purity of the modified oligonucleotides are provided in (Figure S7). Fluorescence measurements were performed at 25 °C using an Edinburgh Instruments FS5 spectrofluorometer to investigate the p47-binding site within the p47A1 aptamer. 1 µM modified aptamer was prepared in a 500-µL volume and loaded in 1-mm path length quartz cuvettes (Starna 110-QS). p47$_{101-266}$ was sequentially titrated to the aptamer solution to yield final protein concentrations of 3, 6, 9, and 12 µM. Sample were excited at 340 nm, and emission spectra were acquired across 355–530 nm. Raw spectral data were processed using Fluoracle software. To assess non-specific fluorescence changes induced by the addition of protein, the UBA domain of p47 (p47$_{1-46}$) was utilized as a negative control under identical conditions (Figure S8).

**NADH coupled ATPase assay**

ATPase activity was measured using a NADH-coupled ATPase assay adapted from previously described methods.[10,11] All reactions contained 25 mM HEPES pH 7.5, 100 mM NaCl, 5 mM $MgCl_2$, 2 mM ATP, 1 mM phosphoenolpyruvate, 0.26 mM NADH, 2.03 U/mL pyruvate kinase, and 5.4 U/mL lactate dehydrogenase. ATP hydrolysis was monitored by measuring the decrease in absorbance at 340 nm at 25 °C. Initial rates were calculated from the linear decrease in NADH absorbance over 5–10 min using an extinction coefficient of 6.22 $mM^{-1}$ $cm^{-1}$. Measurements were performed in triplicate.
